# Bidirectional Peptide Conformation Prediction for MHC Class II using PANDORA

**DOI:** 10.1101/2024.11.04.621877

**Authors:** Daniel T. Rademaker, Farzaneh M. Parizi, Marieke van Vreeswijk, Sanna Eerden, Dario F. Marzella, Li C. Xue

## Abstract

Recent discoveries have transformed our understanding of peptide binding in Major Histocompatibility Complex (MHC) molecules, showing that peptides, for some MHC class II alleles, can bind in a reverse orientation (C-terminus to N-terminus) and can still effectively activate CD4+ T cells. These finding challenges established concepts of immune recognition and suggests new pathways for therapeutic intervention, such as vaccine design.

We present an updated version of PANDORA, which, to the best of our knowledge, is the first tool capable of modeling reversed-bound peptides. Modeling these peptides presents a unique challenge due to the limited structural data available for these orientations in existing databases. PANDORA has overcome this challenge through integrative modeling using algorithmically reversed peptides as templates.

We have validated the new PANDORA feature through a series of experiments, achieving an average backbone binding-core L-RMSD value of 0.63 Å. Notably, it maintained low RMSD values even when using templates from different alleles and peptide sequences. Our results suggest that PANDORA will be an invaluable resource for the immunology community, aiding in the development of targeted immunotherapies and vaccine design.

**Availability:** Source code and data is freely available at https://github.com/X-lab-3D/PANDORA; Contact

## 1 Introduction

MHC class II (MHC II) molecules play a critical role in the immune system by presenting peptides to CD4+ T cells, which are key players in regulating immune responses and influencing the progression of diseases. Traditionally, these peptides were thought to bind to MHC II molecules exclusively in an orientation from the N-terminus to the C-terminus. However, recent discoveries have revealed that peptides can also bind in a reverse orientation, from the C-terminus to the N-terminus for some alleles (Klobuch et al., 2022; Günther et al., 2010; Racle et al., 2023; Schlundt et al., 2012). These discoveries, along with the identification of CD4+ T cells that respond to these reverse-bound peptides (Klobuch et al., 2022), challenge previous assumptions and unveil potential new pathways for therapeutic interventions and a deeper understanding of immune mechanisms.

Understanding the intricate interactions between MHC II molecules and peptides is essential for developing targeted immune therapies and vaccines. The modeling of MHC II peptide conformations has evolved with varied approaches. Most tools, such as pDock (Khan and Ranganathan, 2010) and EpiDock (Atanasova et al., 2013), rely on grid-based docking, while others use homology modeling, including PANDORA (Marzella et al., 2022; Parizi et al., 2023), or more recently, an AlphaFold-based method (Mikhaylov et al., 2024). However, the scarcity of reversed peptides in the Protein Data Bank (Berman et al., 2000; Rademaker et al., 2023) has limited the effectiveness of these methods, highlighting the critical need for more data to drive accurate prediction

Interestingly, Klobuch et al.’s analysis compared canonical and reversed peptides bound to the same MHC molecules, revealing not only conserved anchor residues but also a highly consistent main chain hydrogen bonding pattern across both binding orientations. This consistency suggests that information from canonical peptide X-ray structures can be leveraged to extrapolate the likely configurations of reversed peptides. This approach opens the possibility to use the vast array of existing canonical peptide structures for modeling reverse-bound peptides.

In our current study, we present an update to PANDORA, which now includes a new feature for automatically generating reversed orientation of peptides to be used as templates for homology modeling. This enhancement allows for accurate modeling of reversed-bound peptide configurations. We validated this approach by predicting the conformations of reversed-bound peptides of known MHC II complexes, achieving an average L-RMSD error of 0.63 Å on the peptide binding cores. To the best of our knowledge, PANDORA is the first tool to model reversed-bound peptides, contributing a new capability to the field of immunological research.

## 2 Methods

We created a database of structure templates with reversed peptides by inverting the peptide backbones in the canonical MHC-II complex structures obtained from IMGT (Lefranc et al., 2009), while keeping the side chains untouched.

The preprocessing steps included stripping the PDBs of hydrogen atoms and replacing selenomethionine residues with methionines, which can be properly handled by the Amber14 forcefield (Maier et al., 2015). After generating the initial reversed templates, the peptide residues were renumbered in reverse order, effectively inverting the sequence. Next, we identified the individual planes formed by the carbon (C), nitrogen (N), and oxygen (O) atoms between the alpha carbons (CA) of adjacent amino acids. This step was crucial for preserving the orientation of hydrogen bonds and ensuring that these (N, C, and O) atoms remained in the original plane during the reversal process. Using each plane as a reference, we mirrored the backbone carbonyl (C=O) group with the nitrogen atom across the midpoints between the CAs (see Figures 1A and 1B). After this mirroring process, we reassigned the atoms to the correct amino acids in the PDB file, as the mirroring shifted these atoms to neighboring residues. Finally, reversing the peptide sequence required the removal of the peptide’s C-terminal (COO-) and N-terminal (N-) groups, followed by re-generating these terminal groups to the correct ends of the reversed peptide using PDBFixer (Eastman et al., 2017).

**Figure 1:**
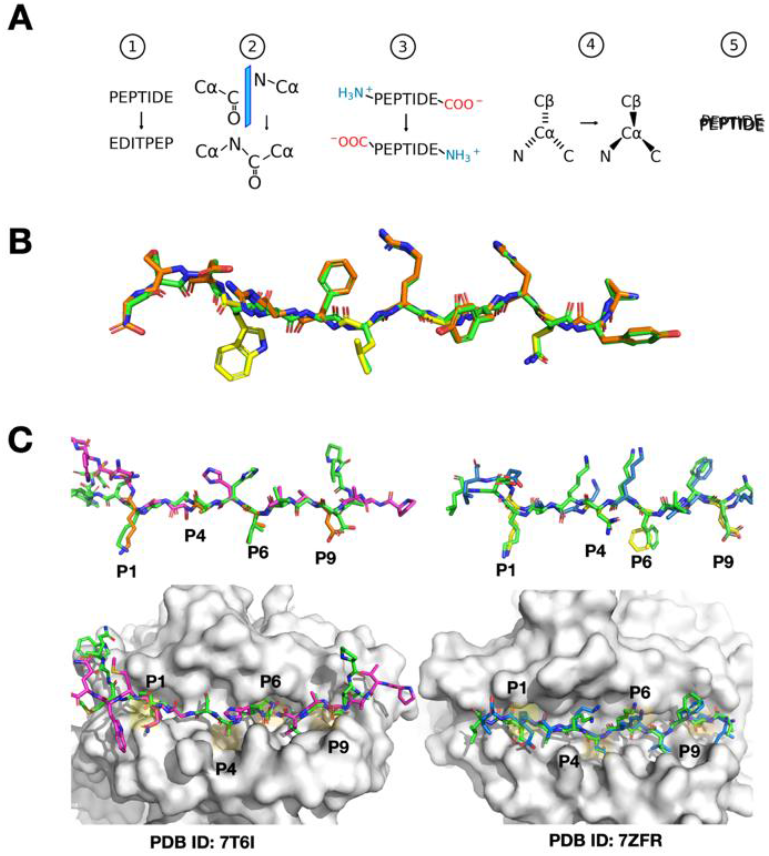
Visual Overview of Peptide Reversal and PANDORA Modeling Predictions. A: Schematic representation of the peptide reversal process in five steps: (1) reversing the sequence numbering and order of the peptide in the PDB file without changing the locations of the residues, (2) mirroring of carbonyl (C=O) and nitrogen (N) groups between the alpha carbons (CA), (3) removing and regenerating terminal groups (NH3+ and COO-) at opposite ends, (4) adjusting CA chirality, and (5) performing molecular dynamics to restore geometry. B: Comparison of the canonical peptide from PDB entry 1AQD (green) and its reversed variant artificially generated by our software (orange, with anchor residues separately highlighted in yellow), illustrating the initial and modified peptide structures. C Left panel: Model of the reversed peptide corresponding to the sequence from PDB entry 7T6I, using 7ZAK_reversed as the template (same allele). The modeled peptide is shown in purple, with main anchor residues highlighted in orange, while the actual X-ray structure of the peptide is depicted in green. This showcases PANDORA’s accurate modeling when using a template from the same allele. The MHC molecule is shown as a white surface, with major binding pockets shaded in yellow. Right panel: Similar to the left panel, but the model corresponds to the sequence from PDB entry 7ZFR, using 3WEX_reversed (Kusano et al., 2014) as the template (different allele). The modeled peptide is shown in blue. This illustrates PANDORA’s ability to generalize to different alleles and accurately predict reversed peptide structures using templates from different alleles.

The rearrangement of atoms resulted in a peptide composed of amino acids in the D-configuration, indicating a reversal of the chirality centers. To achieve the natural L-form, we mirrored the CAs across the plane formed by the N, C, and beta carbon (CB) atoms for each amino acid. Glycines were excluded from this adjustment because they are non-chiral. Similarly, we adjusted the configuration of proline residues by recalculating the CB, CD, and CG side-chain atoms using PDBFixer, resulting in the more common trans isomer.

As the translation of atoms introduced slight distortions, including slight alterations in bond lengths, we performed short molecular dynamics run with OpenMM (Eastman et al., 2017) to restore proper geometry. Before running the MD simulation, we reintroduced hydrogens using PDBFixer and then conducted a 10-step MD run with the Langevin integrator at 300 Kelvin, constraining the MHC complex while allowing only the peptide to move.

After the simulation, we renumbered the anchor residues to ensure consistency with the reversed peptide sequence. For instance, the first anchor residue in the original sequence becomes the last anchor in the reversed sequence, even though it still occupies the same binding pocket in the MHC molecule. The resulting PDB files, containing reversed peptides, were manually inspected, and stored to be used as templates for reverse peptide modeling with PANDORA. All resulting templates are given an ID with the format of PDBID_reversed, e.g., 3WEX_reversed.

## 3 Results

To evaluate the modeling and generalization capabilities of our enhanced PANDORA tool, we conducted two experiments, constrained by the availability of only two structures of reversed peptide-MHC complexes.

The first test involved modeling the reversed MHC-II complex with PDB ID: 7T6I, which includes the DPA102:01-DPB101:01 allele and a reversely-bound peptide PVADAVIHASGKQMWQ. We employed PANDORA, which automatically selected 7ZAK_reversed as the template for modeling, featuring the same alleles and a reversed peptide, DIERVFKGKYKELNK (originally KNLEKYKGKFVREID). This setup allowed us to directly compare the predicted model against the available X-ray structure. We calculated the ligand root-mean-square deviation (L-RMSD) of the binding core residues of the peptide, where L-RMSD measures the average distance between the backbone atoms of the modeled peptide and those in the X-ray structure. The core L-RMSD was 0.746 Å (see Figure 1C), demonstrating PANDORA’s ability to model reversed peptides with high structural accuracy, within the range of the reported core L-RMSD of PANDORA for canonical peptides binding to MHC-II (Parizi et al., 2023).

In our second experiment, designed to assess the generalization capacity of PANDORA, we modeled another MHC-II complex: PDB entry 7ZFR, containing the reversed-bound peptide IEFVFKNKAKEL with the same alleles as in the first experiment (HLA-DPA102:01-HLA-DPB101:01). To challenge PANDORA with a template involving different alleles and a distinct peptide sequence, we deliberately excluded PDB entry 7ZAK from the template selection. PANDORA chose 3WEX_reversed as the template, which contains the reversed peptide FQNFAVTVK (originally KVTVAFNQF) and alleles DPA102:02-DPB105:01. The core-peptide L-RMSD for this second experiment was 0.52 Å (see Figure 1C), aligning with the range of reported core L-RMSD of PANDORA when modeling canonical peptide-MHC II complexes (Parizi et al., 2023). This result suggests that PANDORA can effectively model reversed peptides even when using templates with different alleles and peptide sequences, achieving accuracy comparable to its performance on canonical peptides.

## 4 Discussion and Conclusion

Here we present an updated version of PANDORA, an enhanced tool for modeling MHC II-peptide interactions, now with the added capability to predict reversed peptide bindings. Our results demonstrate that PANDORA can accurately model reversed peptide interactions for specific alleles, as evidenced by the low L-RMSD values obtained in our experiments. Although our testing was limited to two cases due to the scarcity of available reversed peptide-MHC II structures, PANDORA successfully generalized its predictions using both a template from the same allele and one from a different allele. This suggests that PANDORA may have the potential to extend its applicability to model reversed peptides across a broader diversity of polymorphic MHC II molecules.

In conclusion, despite limited structural data, we believe that the updated PANDORA tool will provide the immunology community with a powerful resource for modeling and analyzing reversed peptides in MHC II complexes. This capability may enhance immunotherapy and vaccine design by identifying novel epitopes, opening new avenues for the development of targeted immunotherapies and personalized vaccines. Future work will focus on expanding validation to refine predictions further.

## Acknowledgements

We sincerely thank S. Buchow at Erasmus Medical Center for helpful discussions. FP is supported by the Kika grant (grant number 454). LX and DM are supported by Hanarth Fund and the Hypatia Fellowship from Radboudumc (Rv819.52706). This manuscript benefited from the assistance of ChatGPT, an AI-based language model, in proofreading and refining the language. All final revisions were made by the authors.

## Author Contributions

DR, DM and MV conceptualized the idea of reversing existing canonical-oriented peptide templates. DR designed the algorithms and wrote the initial draft. The experiments were conducted by FP, who also adapted the PANDORA software to enable the modeling of reverse peptides and designed Figure 1. SE and MV extensively revised, tested, and refined the initial code developed by DR. The group leader, LX, provided supervision and guidance throughout the project. DR, FP, and MV are shared first authors. All authors critically reviewed the article.

